# Splenocytes and thymocytes migration patterns between lymphoid organs in pregnancy

**DOI:** 10.1101/2023.08.15.553414

**Authors:** Gabriela T. Cruz-Cureño, Marina Ch. Rosales-Tarteaut, Lourdes A. Arriaga-Pizano, Luvia E. Sánchez-Torres, Denisse Castro-Eguiluz, Jessica L. Prieto-Chávez, Rodolfo Pastelin-Palacios, Ana Flisser, Arturo Cérbulo-Vázquez

## Abstract

**Background:** Cell migration is essential for the immune system, and frequently is analyzing in adult non-pregnant animals, and poorly explored in pregnancy, however, a physiologic increase of size in the spleen and periaortic lymph nodes had been reported in pregnant mice.

**Methods:** Using a mouse model, we transferred PKH26-stained thymocytes and splenocytes from pregnant or non-pregnant animals to receptor mice in the presence or absence of pregnancy. Percentage and Mean Fluorescence Intensity were calculated by Flow cytometry in pregnant or non-pregnant mice. Non-parametric ANOVA analysis was performed.

**Results:** We detected that the percentage of PKH26+ thymocytes into the spleen, lymph nodes, and peripheral blood is higher in females than in males. Our results showed a similar frequency of thymocytes and splenocytes from pregnant and non-pregnant located into receptor lymphoid organs. Also, the location of marked cells was similar during the perinatal period.

**Conclusions:** The mobility of thymocytes and splenocytes in pregnant and non-pregnant mice is similar, therefore we suggest that the larger size of the spleen and periaortic lymph nodes noted previously in pregnant mice, could be the result of retention of leukocytes in the secondary lymphoid organs.

## Background

Cell trafficking is an essential function of the immune system. Cell migration patterns had been recognized for leukocytes egressing from bone marrow and thymus (primary lymphoid tissues) into the blood and afterwards to lymphnodes and spleen (secondary lymphoid tissues, SLO) (1, 2). Also, thymocyte migration within the thymus has been detected as critical for positive and negative selection (3). While, naïve T and B cells that egress from thymus or bone marrow could recognize antigen in the SLO, and then proceed to activation and proliferation (4). An alternative behavior, depending on antigen encounter, is for naïve lymphocytes to egress from SLO, and return to the blood through the thoracic duct and recirculate (5). In addition, memory cells follow a similar migration pattern as naïve cells; in contrast, effector cells home to non-lymphoid tissue and express their function (6, 7).

Also, time is important to analyzed the pattern of cell migration, mature T cells that egress from the thymus only spend minutes in the blood (8, 9), and then they arrives to the SLO (4, 10). Lymphocytes remain in the SLO for 8-12 hours (T cells) or 24 hours (B cells) searching for antigen, if they encounter their antigen they remain in the SLO, and become activated and proliferate; if no antigen is encountered, they leave the SLO and circulate in blood (8). Pattern of cell migration also change thorough the day, and has been reported that the circadian clock controls the leucocyte trafficking (11), were the peak of leukocyte circulation in rodents is during the day (12).

Immune cells’ migration patterns haven’t been characterized in pregnancy, where the immune system express a high regulated state of immunotolerance. Reports have described an increase of leukocyte migration to the paraaortic lymphnodes in pregnant mice, with hypertrophy between 14 and 16 days (13). Also, a reduction in the size of the thymus and an increase in the size of the spleen have been reported previously in pregnancy (14), these changes are observed both in allogeneic and syngeneic systems (15, 16, 17, 18). Similar to the spleen, the proportion and the absolute number of mononuclear cells in peripheral blood show an increase during pregnancy (19).

In this study we aimed to analyze whether cell migration to lymphoid organs depends entirely on the cell or if it is conditioned by the pregnancy environment. We used a stained cell transfer model in mice. Thymocytes or splenocytes from pregnant or non-pregnant animals were isolated, marked and transferred to pregnant or non-pregnant pairs. We report the proportion and absolute numbers of cells in the thymus, spleen, lymphnodes and blood. Female mice showed a higher percentage of marked cells to lymphoid organs than male mice. The proportion of splenocytes detected in the spleen were higher than the thymocytes detected in the spleen. Our results suggest that the cell condition alone supports a differential location to the lymphoid organs, suggesting that the migration program is expressed in leukocytes and minor affected by the pregnant environment.

## Methods

An experimental, prospective, and analytical study was conducted to describe if the frequency of marked leucocytes in lymphoid organs is similar in absence and presence of pregnancy.

### Mice

Female and male BALB/c mice (3 mice per experimental group), 6-8 weeks of age, were obtained from Bioterio the Animal Production and Experimentation Unit (UPEAL-B), UAM-Xochimilco. The management of biological samples was carried out according to the Biosafety Manuals of the Faculty of Medicine of the UNAM. Mice were kept in acrylic boxes (19×29×12cm), under constant temperature conditions (23ºC) with standard 12 h light/dark intervals. Food and water were supplied on demand. To analyze homogeneous groups of mice, the hormonal cycle was tested and animals in the estrous cycle were used. Briefly, the vaginal sample was taken every 24 hours (four consecutive days) and stained with 0.1% crystal violet. The colpocytological exams were observed under an optical microscope, showing a physiologic cyclicity.

### Collection of peripheral blood

Sevoflurane was used as an inhalation anesthetic, the lack of reflexes, relaxation, and regular breathing was checked before starting the procedure. Mice were placed supine and peripheral blood was extracted from mice by cardiac puncture. Briefly, locating the xiphoid bone, a lateral puncture with an angle of 30º was performed, and exerting gentile negative pressure with the syringe (1mL volume/25G needle) blood emerges. Then mice were sacrificed by cervical dislocation. The blood was collected in microtainer tubes (BD, catalog. 363706 NJ, USA). Subsequently, and using NH_4_Cl solution (150 mM ammonium chloride, 10 mM KHCO3, and 1mM EDTA) the erythrocyte lysis was performed. The sample was washed with PBS (Sigma-Aldrich, catalog P3813 St. Louis, MO) at 4°C, and centrifuged at 1500 rpm for 10 minutes. The suspension was decanted, and the cell button was resuspended in 100 μL. Then using a Neubauer chamber cell count and viability were check by the trypan blue exclusion method.

### Cell suspension from lymphoid organs

Pregnant (P) and non-pregnant (NP) adult mice of 6-8 weeks under anesthetic, were sacrificed by cervical dislocation. Then, the thymus, spleen, axillary and inguinal lymph nodes were dissected. Cell suspensions were obtained using a fine mesh and washed by centrifugation (PBS at 4°C, at 1,200 rpm for 10 minutes). The Samples were decanted, filtered, and resuspended in a volume of 500 μL. Cell count and viability were evaluated by the trypan blue exclusion method.

### PKH-26 mark of cells

Five million cells were marked using the PKH-26 Red Fluorescent Cell Linker Kits (Sigma-Aldrich, catalog PKH26GL St. Louis, MO). Following the manufacture instructions, cells were resuspended in diluent C (Sigma-Aldrich, catalog CGLDIL St. Louis, MO), and mixed with PKH26 ethanolic dye solution (Sigma-Aldrich, catalog P9691 St. Louis, MO) and incubating 5 minutes at 37°C. After incubation time the reaction was stopped with PBS (Sigma-Aldrich, catalog P3813 St. Louis, MO), tubes centrifuged, and cells resuspended in 200 μL of PBS. Cell count and viability were checked as previously. Data acquisition and analysis were performed on a FACSCanto® (BD Biosciences) using the Infinicyt™ software (Cytognos Euroflow®) for data analysis. Fig. 2a-d show the plots for PKH-26+ thymocytes identification in receptor lymphoid organs and peripheral blood, while Fig. 2e-h show the plots for PKH-26+ splenocytes in receptor organs and blood.

### Transfer of cells

Four cell transfer systems were analyzed: **NP→NP, NP→P, P→NP**, and **P→P**. Local heat was used to dilatate the caudal vein, then the tail was clamped. Using insulin syringes with a 13 mm long (1/2”) and 27G thick needle (BD, catalog 326716 NJ, USA), 5×10^6^ of PKH26+ cells in a volume of 200 μL were transferred at the 15-16 days post coitus (dpc), or in the estrous phase if mice were non-pregnant. All cell transfers were performed early in the day (between 9:00 and 11:00 h).

### Count of PKH26+cells in receptor lymphoid organs

Two days after the cell transfer, mice were sacrificed and lymphoid organs isolated. One million cells per 100 μL were obtained from the spleen, thymus, lymph node, peripheral blood or placenta, and placed into cytometry tubes (Falcon, catalog 352008 MA, USA). Then following manufacturer instructions 300 μL of FACS Lysing Solution was added (BD, catalog 349202 San Jose, CA), and incubated in dark for 10 minutes. Then, one mL of PBS was added and cells were centrifuged at 3,000 rpm for 5 minutes, samples were decanted and resuspend in 100 μL of PBS. Data acquisition was performed on the FACSCanto flow cytometer (BD Biosciences, USA), and using Infinicyt cytognos euroflow software for analysis (Cytognos SL, Salamanca, Spain). The strategy to identify PHK26+ cells used dot plots graphs (Fig. 1).

**Figure 1.**
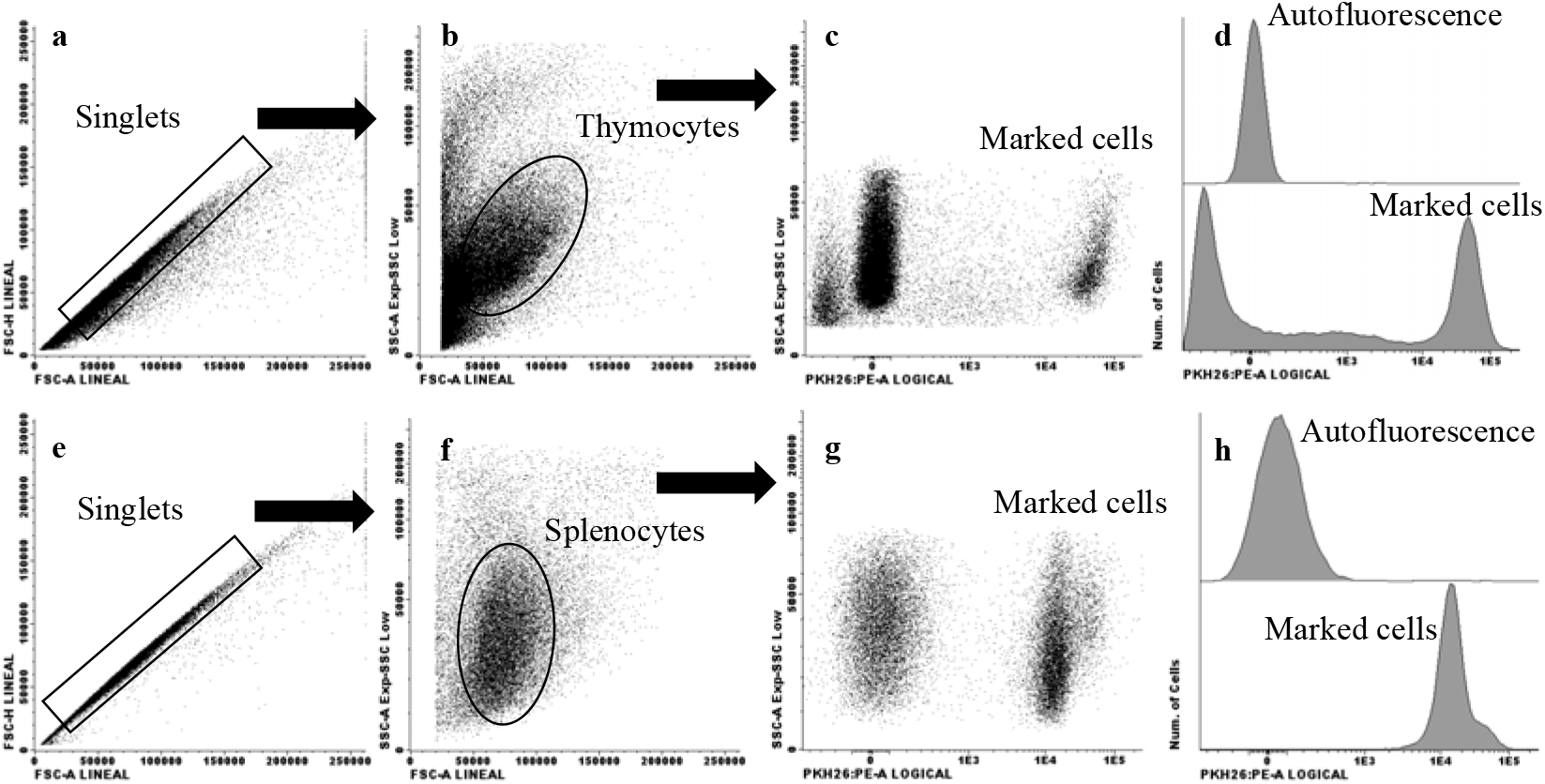
Thymocytes or splenocytes after labeling with PKH-26. Cell suspensions were obtained from the thymus or spleen, and labeled with PKH-26 as in methods (1a-h). Images show the total cells acquired per organ of non-pregnant mice (n=3). Top histogram shows unmarked cells, and bottom histogram show PKH-26+ in thymocytes (Fig. 1d) and splenocytes (Fig. 1h).

### Statistics

Analysis was performed using GraphPad Prism version 6 software (GraphPad software Inc. La Jolla, California, USA). Non-parametric Kruskal Wallis test with Dunn post-test multiple comparisons test was used to evaluate differences in percentages of cells, and in the Mean Fluorescence Intensity (MFI) among the groups. A p<0.05 was considered statistically significant. No power calculation was performed.

## Results

### Percentage of cells PKH26 marked and viability

Thymocytes or splenocytes were labeled for PKH26 and up to 95% cell viability was obtained. Single thymocytes and splenocytes were analyzed, as shown (FSC-A *vs*. FSC-H plot) in Fig. 1a and 1e respectively. A classical pattern of size *vs*. granularity was obtained for thymocytes (Fig. 1b) and splenocytes (Fig. 1f). PKH-26 *vs*. granularity dot plots show PKH-26 positive thymocytes (Fig. 1c) and splenocytes (Fig. 1g), up to 84% of thymocytes were PKH-26+, and around 90% of splenocytes were marked. Fig. 1d and 1h show autofluorescence and marked thymocytes and splenocytes, respectively. The presence of marked thymocytes (Fig. 2a-2d) or splenocytes (Fig 2e-2h) transferred from NP mice to NP mice in thymus, spleen, blood and lymphnodes is described.

**Figure 2.**
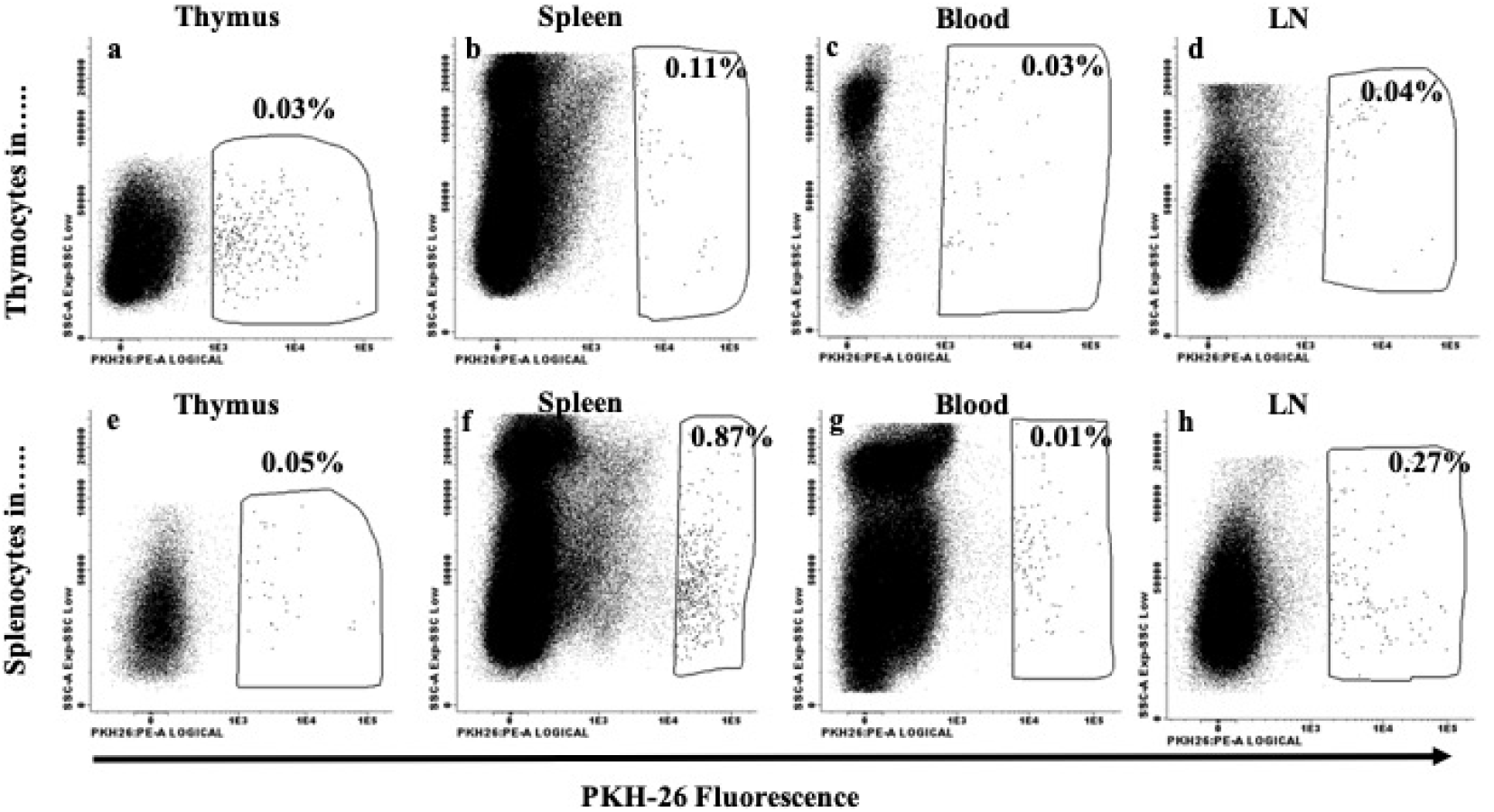
PKH-26+ thymocytes or splenocytes in receptor organs. Cell suspensions were obtained from the thymus or spleen, and labeled with PKH-26 as in Fig 1. PKH-26+ cells were identified two days after in receptor lymphoid organs and peripheral blood (2a-h). Images show the total cells acquired per organ of non-pregnant mice (n=3). Gated cell shows PKH-26+ cells in lymphoid organs and peripheral blood (Fig 2a-h).

### The percentage of PKH-26+ cells were higher in female than in male mice

PKH-26+ cells were transferred and after 48 hours receptor lymphoid organs were collected. Marked cells were quantified by flow cytometry in female and male mice. In Figure 3A we analyzed the cell migration dynamics comparing female and male receptor mice. In most organs we observed a higher frequency of PKH-26+ cells in female samples than in males (n=4), except for male thymus. Also, the percentage of PKH-26+ splenocytes in female spleen was higher than in the male spleen (p=0.039), lymphnodes (p=0.039) and peripheral blood (p=0.039).

**Figure 3.**
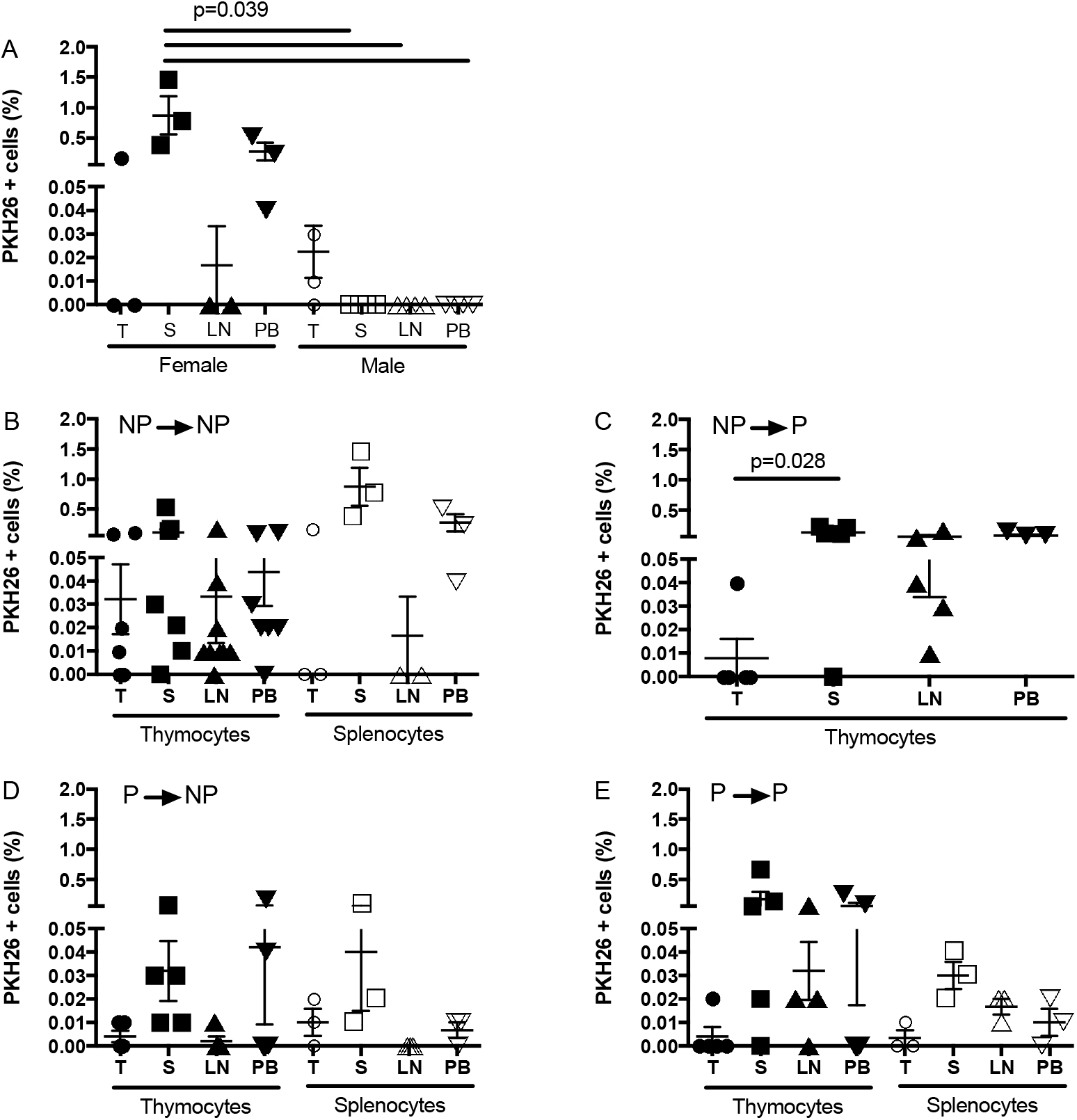
Percentage of PKH-26+ cells in receptor lymphoid organs and peripheral blood. Thymocytes or splenocytes were PKH-26 marked as in methods. Five million cells were transferred to the receptor mouse and counted two days after by flow cytometry. A) PKH-26+ thymocytes were counted in lymphoid organs in female and male mice. B-E) PKH-26+ thymocytes or splenocytes were quantified in lymphoid organs in the following groups: **NP→NP, NP→P, P→NP** and **P→P**. Data represented as mean±SEM. Kruskal-Wallis test, Dunn’s multiple comparisons test, IC 95%. The significance value was p<0.05. T: thymus; S: spleen; LN: lymphnodes; PB: peripheral blood. Filled symbols: Thymocytes. Open symbols: Splenocytes. Each symbol shows a mouse.

### PKH-26+ location to lymphoid tissues in female mice with and without pregnancy

Figures 1 and 2 shows the strategy to label cell with PKH-26 and count those cells in receptor tissues, in Fig. 3 we show the percentage of PKH-26+ cells for each mouse and every condition with NP or P mice. The percentage of PKH-26+ thymocytes and splenocytes from **NP→NP** mice is shown in Fig. 3B. No statistical difference was found between PKH-26+ thymocytes and splenocytes in receptor lymphoid organs. In the case of **NP→P** group (Fig. 3C), we observed a statistical difference between PKH-26+ thymocytes in spleen vs. in thymus (p=0.028), and no statistical difference was noted among the other lymphoid organs. For the **P→NP** and **P→P** groups (Fig. 3 D and E, respectively) no statistical difference was observed between PKH-26+ thymocytes and splenocytes, also no difference was described among receptor lymphoid organs. We observed a significant difference between the PKH-26+ thymocytes in lymphnodes for **NP→P *vs*. P→NP**, with a higher frequency of cells in the **NP→P** group (Fig. 3C vs D).

Figure 4 shows the absolute number of PKH-26+ thymocytes or splenocytes quantified 48 hours after the transfer. The highest number of cells were detected in the **NP→NP** condition, however this was not statistically different compared to the other groups (**NP→P, P→NP** or **P→P**). In general, the number of PKH-26+ splenocytes detected in the lymphoid organs was higher than the number of PKH-26+ thymocytes; however, no statistically significant differences were observed. The number of PKH-26+ splenocytes was higher in the spleen compared to LN in the **NP→NP** group (p=0.047).

**Figure 4.**
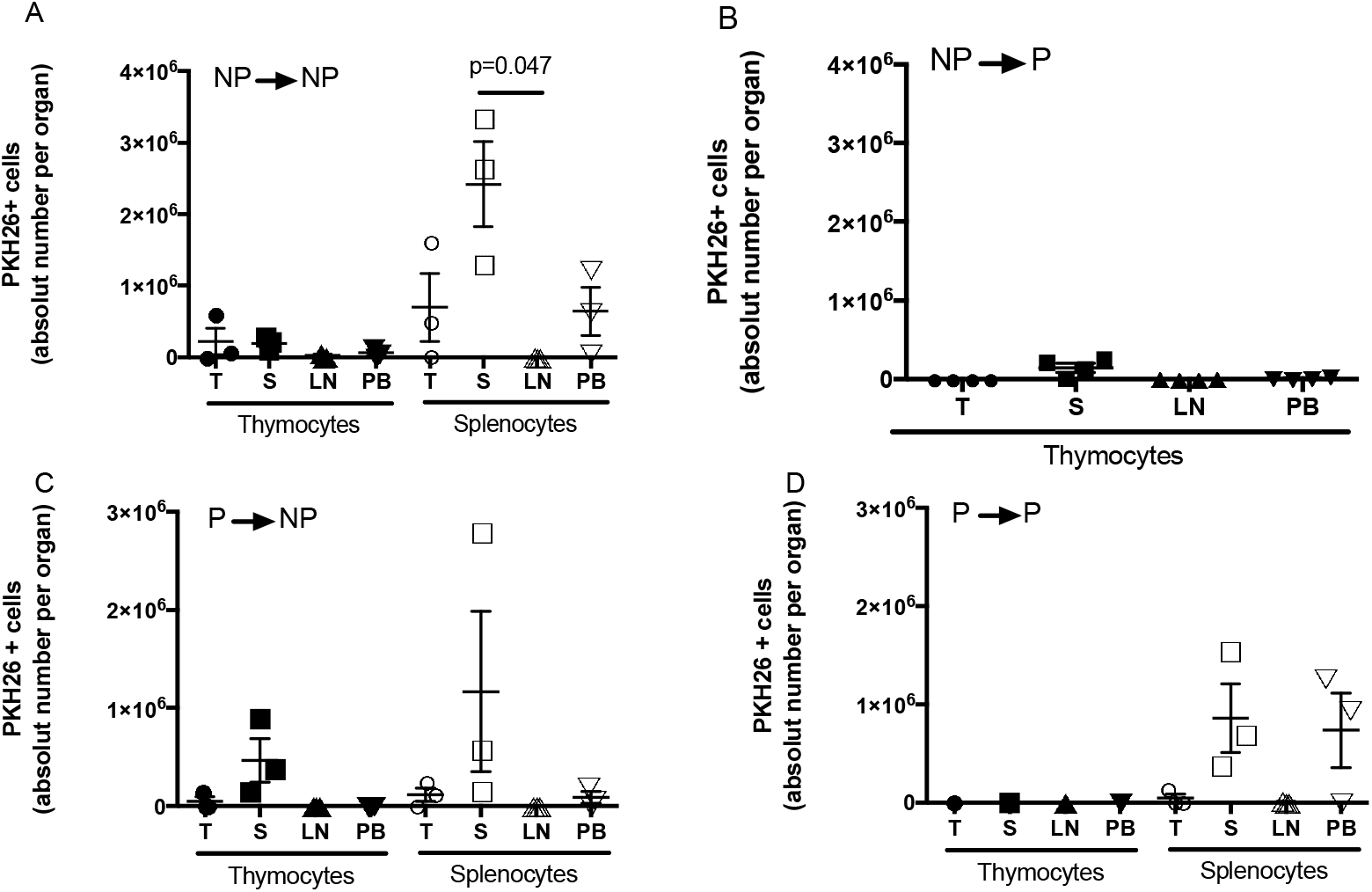
Absolut number of PKH-26+ cells in receptor lymphoid organs and peripheral blood. Thymocytes or splenocytes were PKH-26 marked as in methods. Five million cells were transferred to the receptor mouse and quantified after 48 hours by flow cytometry. Data represents mean±SEM. Kruskal-Wallis test, Dunn’s multiple comparisons test, IC 95%. The significance value was p<0.05. T: thymus; S: spleen; LN: lymphnodes; PB: peripheral blood. Filled symbols: Thymocytes. Open symbols: Splenocytes. Each symbol shows a mouse.

Finally, to explore if PKH-26+ cells express a differential location to lymphoid organs around the moment of birth, PKH-26+ thymocytes or splenocytes were transferred from Pregnant (15-16 dpc) to PostPregnant (PP) mice (1-3 days after birth). A similar frequency of thymocytes and splenocytes in lymphoid organs was observed in the PP receptor mice, compared to NP or P mice (Suppl. Table 1).

**Suppl. Table 1.** Presence of transfer PKH26+ cells in lymphoid organs or blood in the perinatal period.

| Donor → | Thymus |  |  |  |  | Spleen |  |  |  |  |
| --- | --- | --- | --- | --- | --- | --- | --- | --- | --- | --- |
| Receptor → | T<br>(n=3) | S<br>(n=3) | LN<br>(n=3) | PB<br>(n=3) | *p* | T<br>(n=3) | S<br>(n=3) | LN<br>(n=3) | PB<br>(n=3) | *P* |
| P → NP | 0.003±<br>0.01 | 0.02±<br>0.01 | 0 | 0 | ns | 0.01±<br>0.01 | 0.04±<br>0.04 | 0 | 0.01±<br>0.01 | ns |
| P → P | 0.007±<br>0.01 | 0.27±<br>0.34 | 0.04±<br>0.02 | 0.08±<br>0.14 | ns | 0.003±<br>0.006 | 0.03±<br>0.01 | 0.02±<br>0.01 | 0.01±<br>0.01 | ns |
| P → PP | 0.01±<br>0.01 | 0.12±<br>0.10 | 0.06±<br>0.05 | 0.14±<br>0.16 | ns | ND | ND | ND | ND | -- |
| *p'* | ns | ns | ns | ns |  | ns | ns | ns | ns |  |

Thymus (T), Spleen (S), Lymphnode (LN), Peripheral Blood (PB). Data represented as mean±SD. Kruskal-Wallis test, Dunn's multiple comparisons test, IC 95%. The significance value was $p < 0.05$ . ns: not statistically significant; ND: not determined.

## Discussion

Several physiological changes take place during pregnancy, among these changes, an increase in plasma and blood cells has been thoroughly described (20, 21). We asked if during this physiological leukocytosis the leukocyte mobility could follow a different migration pattern compared to the absence of pregnancy. Using an experimental model, we addressed this problem. We report the proportion of PKH26+ thymocytes and splenocytes located in different lymphoid tissues in the presence or absence of pregnancy.

The traffic of leukocytes among lymphoid organs increases in some phases of pregnancy, it has been reported that the traffic of monocytes towards the uterus is necessary for the correct development of the initial phases of labor by increasing uterine contractility (22). In physiologic conditions without pregnancy, classic cell migration pattern goes from primary to secondary lymphoid tissue, and then to non-lymphoid tissue where they do an effector function (23). Also, recently it has been determined that primary lymphoid organs are also in the pathway of lymphocyte trafficking (24, 25, 26).

First, when we compared the proportion of PKH26+ cells from female mice and transferred to female and male mice, we observed that female mice have had a higher proportion of transfer cells in lymphoid tissue than male mice, indicating that the cell migration is more frequent in females than males. Also, it could indicate that our observations in pregnant and non-pregnant mice are valid because males showed a very low number of PKH-26+ cells, in contrast, cells persisted (at least for two days) in females in all the tissues analyzed.

The number of marked cells in a particular tissue depends on the migration rate into the tissue, the proliferation *in situ*, cell death, or the outflow from the tissue, also, pregnancy is a unique condition where the change in the cellularity and size of lymphoid organs have been reported, we hypothesized that the cell itself or the pregnancy environment could act to regulate the migration of leukocytes in mice. In agreement with a previous report (14), we observed that the absolute number of cells increase in the spleen of pregnant mice. Also, our results showed that the number of PKH26+ splenocytes is higher in the spleen of pregnant mice, however we do not know if this is because a higher cell proliferation or a lower outflow cells from the tissue. More studies are necessary to clarify this question.

Our results showed that the proportion of PKH26+ cells in lymphoid organs were similar between pregnant and non-pregnant condition. We observed that PKH26+ splenocytes were located mainly in the spleen and to a lesser extent to lymph nodes. The percentage of labeled thymocytes found in the spleen and lymph nodes was low, which would indicate that these tissues are populated by mature resting lymphocytes. Also, in agreement with a previous report (26), the number of PKH-26+ thymocytes detected in the thymus was low in most cases indicating that thymocytes can goes back to the thymus, but the pregnancy do not alter this pattern. Since the proportion of PKH26+ thymocytes or splenocytes is higher in organs than in the peripheral blood, we believe that the number of PKH26+ cells reflect the proportion of cell real located in the organ. More analysis must be performed to solve this question. Finally, our results suggest that the mobility of thymocytes and splenocytes in the receptor mice is similar between pregnancy and near the time after birth. Also, a percentage of PKH-26+ cells were detected in the placenta of receptor pregnant mice (data do not show), and the number was similar to the lymphoid organs analyzed, suggesting that the placenta could support certain immune vigilance and interchange between mother and fetus. The percentage of microchimerism in humans detected with molecular biology or histologic test is similar to the percentage of cells that we detected (27, 28, 29), this open the option to explore aspects of microchimerism in a mice model, doing experiments that are not possible in humans for ethical reasons.

## Conclusions

Splenocytes and thymocytes have similar migration patterns to lymphoid organs irrespective of pregnancy condition, but female receptors persistently retain transferred cells.

## Abbreviations

SLO: Secondary Lymphoid Organs
NP: Non-Pregnant
P: Pregnant
dpc: days post coitus
MFI: Mean Fluorescence Intensity
T: Thymus
S: Spleen
LN: Lymphnode
PB: Peripheral Blood
PP: PostPregnant

## Declarations

### Ethical considerations

The experiments were performed according the guidelines for the use of laboratory animals in the Escuela Nacional de Ciencias Biológicas del Instituto Politécnico Nacional. The study was approved by the Animal Ethics Committee of the Escuela Nacional de Ciencias Biológicas del Instituto Politécnico Nacional, Mexico (ZOO-025-2018).

### Consent for publication

Not applicable

### Availability of data and materials

The datasets used and/or analyzed during the current study are available from the corresponding author on reasonable request.

### Authors’ contributions

ACV conceived and designed the study, and contributed to the analysis of data. GCC and MRT write the first version of the report, and LAP, LST, RPP, AF contributed to the critical revision of the report. GCC, MRT, LAP, LST, JPCh, RPP, and AF, contributed to data acquisition, data analysis or data interpretation. All authors reviewed the final version. ACV reviewed and approved the final version.

## Acknowledgements

The authors thank the staff at Bioterio UPEAL-Xochimilco and Bioterio del Departamento de Inmunología, ENCB-IPN, HGM, Fac Med UNAM. MSc Gabriela T. Cruz-Cureño and MSc Marina Ch. Rosales-Tarteaut got CONACyT scholarship (619436 and 58442 respectively).

## Authors’ information (optional)

Gabriela T. Cruz-Cureño, Marina Ch. Rosales-Tarteaut, Luvia E. Sánchez-Torres: Escuela Nacional de Ciencias Biológicas, IPN. Prolongación Carpio s/n esq. Plan de Ayala, Plutarco Elías Calles, CP 11340. Ciudad de México

Lourdes A. Arriaga-Pizano, Jessica L. Prieto-Chávez: Unidad de Investigación Medica en Inmunoquímica, Centro de Instrumentos, Hospital de Especialidades, CMN Siglo XXI, Av. Cuahutemoc 330, Doctores. CP 06725. Ciudad de México

Rodolfo Pastelin-Palacios: Facultad de Química, Universidad Nacional Autónoma de México. Ciudad Universitaria. Circuito interior. Av. Universidad 3000. CP 04510. Ciudad de México. Mexico

Ana Flisser: Departamento de Microbiología y Parasitología, Facultad de Medicina, Universidad Nacional Autónoma de México. Ciudad Universitaria. Circuito interior. Av. Universidad 3000. CP 04510. Ciudad de México. Mexico

Arturo Cérbulo-Vázquez: Hospital General de México “Dr. Eduardo Liceaga”. Medicina Genómica. Ciudad de México. Dr. Balmis 148. Doctores. Alcaldía Cuauhtémoc. CP 06726. Ciudad de México. México.

